# Biofilm-associated *Mycobacterium abscessus* cells have altered antibiotic tolerance and surface glycolipids in Artificial Cystic Fibrosis Sputum Media

**DOI:** 10.1101/479923

**Authors:** Augusto Cesar Hunt-Serracin, Brian J. Parks, Joseph Boll, Cara Boutte

## Abstract

*Mycobacterium abscessus* (*Mab*) is a biofilm-forming, multi-drug resistant, non-tuberculous mycobacterial (NTM) pathogen increasingly found in Cystic Fibrosis patients. Antibiotic treatment for these infections is often unsuccessful, partly due to *Mab*’s high intrinsic antibiotic resistance. It is not clear whether antibiotic tolerance caused by biofilm formation also contributes to poor treatment outcomes. We studied the surface glycolipids and antibiotic tolerance of *Mab* biofilms grown in Artificial Cystic Fibrosis Sputum (ACFS) media in order to determine how they are affected by nutrient conditions that mimic infection. We found that *Mab* displays more of the virulence lipid trehalose dimycolate when grown in ACFS compared to standard lab media. In ACFS media, biofilm-associated cells are more antibiotic tolerant than planktonic cells in the same well. This contrasts with standard lab medias, where biofilm and planktonic cells are both highly antibiotic tolerant. These results indicate that *Mab* cell physiology in biofilms depends on environmental factors, and that nutrient conditions found within Cystic Fibrosis infections could contribute to both increased virulence and antibiotic tolerance.

## Introduction

Infections caused by Non-Tuberculous Mycobacteria (NTM) are on the rise across the globe (1-3). *Mycobacterium abscessus* (*Mab*) is a highly antibiotic-resistant NTM that causes soft tissue infections and is an increasingly common respiratory pathogen in Cystic Fibrosis (CF) patients (4-6). While most NTM infections are believed to be contracted through environmental exposure, *Mab* has recently been shown to be transmitted between CF patients, probably by fomites (7). Unfortunately, *Mab* infections in CF patients are very difficult to treat, and there are no standardized treatment regimens. Treatment usually involves some combination of clarithromycin, amikacin, imipenem, cefoxitin and/ or linezolid(8). Infection clearance after up to 3 years of antibiotic treatment is achieved in only 10-55% of patients (8). This level of treatment failure is likely partly due to intrinsic antibiotic resistance (9-11); however, antibiotic tolerance may also play a role (12), though the environmental conditions that induce tolerance in this organism are poorly understood.

The pathophysiology of *Mab* infection in the CF lung has scarcely been studied, but one pathology study of explanted lungs from CF patients showed *Mab* aggregates that appear to be forming a biofilm around the alveoli (13). Biofilms are bacterial communities that in many species are held together by exopolysaccharides (14). Mycobacterial biofilms appear to depend on surface glycolipids (15) and free mycolic acids (16, 17), though many questions remain about the composition of the matrix in mycobacterial biofilms. Because bacteria growing in biofilms are notoriously tolerant to antibiotics (18), this mode of growth during infection may contribute to the treatment recalcitrance of *Mab*. While we currently lack a comprehensive model of the chemical and genetic basis of *Mab* biofilm formation, it is clear that surface glycolipids are important. Glycopeptidolipids, which are found on the outer leaflet of the mycobacterial outer membrane, have been shown to affect biofilm structure (19) and are thought to modulate the course of infection (20, 21). Trehalose dimycolate (TDM), which is known to contribute to virulence in *Mtb* (22) are also found on the surface of some *Mab* strains, though its contribution to biofilm formation and virulence in *Mab* has not been studied.

*Mab* has two genetic isoforms: the wild-type “smooth” strains, which produce abundant GPLs and minimal TDM, and “rough” mutants, which have genetic lesions in the GPL loci and display little or no GPLs but sometimes have higher levels of TDM (23, 24). The smooth strains form more robust biofilms and it is thought that this form could contribute to colonization at infection sites (24), which is likely promoted by GPLs masking other cell surface molecules that activate innate immunity (21). The rough mutants cause more inflammation (21), are more virulent, and are thought to promote more tissue invasion (24). Rough mutants are more frequently isolated after a persistent infection has been established for a long period, in both CF and non-CF patients (25, 26).

The smooth and rough genetic variants represent two phenotypic extremes of *Mab* physiology. However, the rough variants only form in some infections, and it is thought that most infections are established by smooth strains (24). Smooth strains are known to form biofilms *in vitro* (12, 19) and are thought to also form biofilms during infection (13). Antibiotic-recalcitrant *Mab* infections are likely to involve mature biofilms growing in complex nutrient environments. However, *in vitro Mab* biofilm studies have been done on immature biofilms in simple nutrient environments (12, 19): in these conditions there are modest differences in antibiotic tolerance between biofilm-associated and planktonic cells (19). Thus, it is not clear to what extent biofilms contribute to antibiotic tolerance during infection.

In this work, we sought to determine how cell surface physiology, biofilm formation and antibiotic tolerance are affected by media conditions in *Mab* ATCC19977. We find that the cell surface, as measured by fluorescent staining, is different in *Mab* in Artificial Cystic Fibrosis Sputum (ACFS) media compared to either minimal HdB media or in the standard mycobacterial growth media, 7H9. In addition, both smooth and rough strains display more TDM in the ACFS media. When we compare biofilm and planktonic cells, we find that both populations are quite antibiotic tolerant in HdB and 7H9, while the planktonic cells are more susceptible to antibiotics in ACFS media.

## Results

To determine how different nutrient conditions affect basic cell physiology, we measured growth rate and microscopically examined *Mab* in three different media. *Mab* grows fastest in 7H9 media (Fig 1AB), even though the Artificial Cystic Fibrosis Sputum (ACFS) media is richer (27). *Mab* in ACFS media appears to die off more in stationary phase, implying that the nutrient richness may not prepare cells for certain stresses (Fig. 1A). We imaged cells using phase microscopy and observed that cell lengths were longer in ACFS media (Fig. 1CD). We stained log. phase cells in each media with both NADA, a 4-chloro-7-nitrobenzofurazan-conjugated D-alanine (28), which is integrated into peptidoglycan, and FM4-64, which stains the inner membrane (29). Although growth rates were more similar in HdB and ACFS media, cell surface staining was more similar in HdB and 7H9. The NADA staining in Hdb and 7H9 was consistent with studies performed in other mycobacterial cells: brighter staining at the septa and poles where new peptidoglycan is inserted (30). FM4-64 stained most cells in these media types dimly, but a subset stained more brightly. Staining of cells in ACFS media was more homogeneous (Fig. 1DE). We observed in the phase images that cells growing in ACFS appear to be encased in translucent sheaths, which could alter access of the stains to the cell (Fig. 1D). These data show that *Mab* is physiologically different in typical lab media than in media that mimics the nutrient conditions in the lungs of Cystic Fibrosis patients.

**Figure 1.**
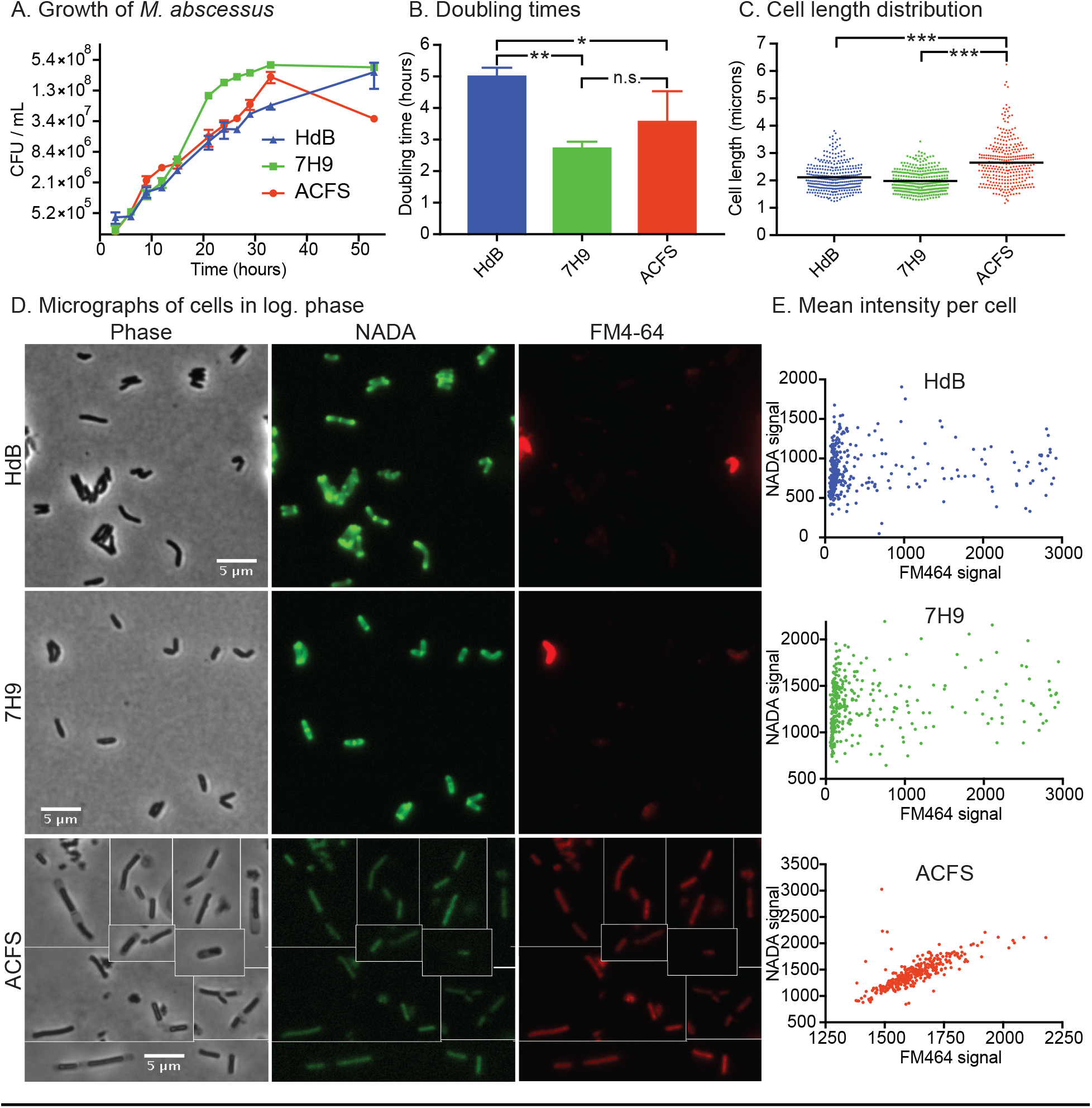
The physiology of *Mab* planktonic cells varies according to media type. A) Colony forming units over time in the three growth media on a 2-log scale. B) Maximum doubling times during log. phase growth, calculated from data in (A). P values calculated by one-way ANOVA with Tukey correction. Hdb vs. 7H9, P=0.0059; Hdb vs.ACFS, P=0.0459. C) Quantification of cell lengths of *Mab* cells in log. phase, in the three media types. At least 300 cells from two biological replicates were imaged by phase microscopy and cell lengths were calculated using MicrobeJ. ACFS vs. HdB, P<0.0001; ACFS vs. 7H9, P<0.0001. D) Micrographs of log. phase cells stained with the fluorescent D-amino acid NADA and the membrane stain FM4-64. E) Quantification of the average fluorescent intensity for at least 300 cells from two biological replicates, stained with NADA and FM4-64, as in (D).

In order to study the physiology of biofilms, we sought to develop a method to separate biofilm and planktonic cells from the same culture well: this allows us to control internally for nutrient depletion in the media, which also impacts cell physiology and antibiotic tolerance. We incubated standing cultures in tissue culture-treated plates. Biofilms form at the bottom of each well, while the planktonic cells remained in the media in the upper half of the well. We pipetted off all the media on the top of each well to isolate planktonic cells, then resuspended the surface-associated cells on the bottom of each well in 7H9 with Tween80, a non-ionic detergent, which broke apart cell clumps. To determine whether 7H9 + Tween80 was sufficient to break apart clumps or whether sonication was necessary, we compared the colony forming units (CFU) produced from biofilm resuspensions with and without sonication (Fig. S1). Sonication did not change the CFU counts, so we conclude that resuspension in 7H9 + Tween80 alone is sufficient to break apart smooth *Mab* biofilms. Therefore, we used that method for biofilm enumeration in subsequent experiments.

We next analyzed how the three media types affect biofilm development and morphology. We grew *Mab* in the three different media types over time and monitored biofilm development by CFU assay and photography. Biofilms growing in all media types reach maximal cell density by day 6 (Fig. 2). Biofilms in ACFS form three-dimensional plumes in standing culture, while those in HdB or 7H9 are flat and almost crystalline. We did not observe pellicles in the media conditions we tested using smooth *Mab* strains (Fig. 2). From these data we conclude that biofilm morphology was affected by media condition.

**Figure 2.**
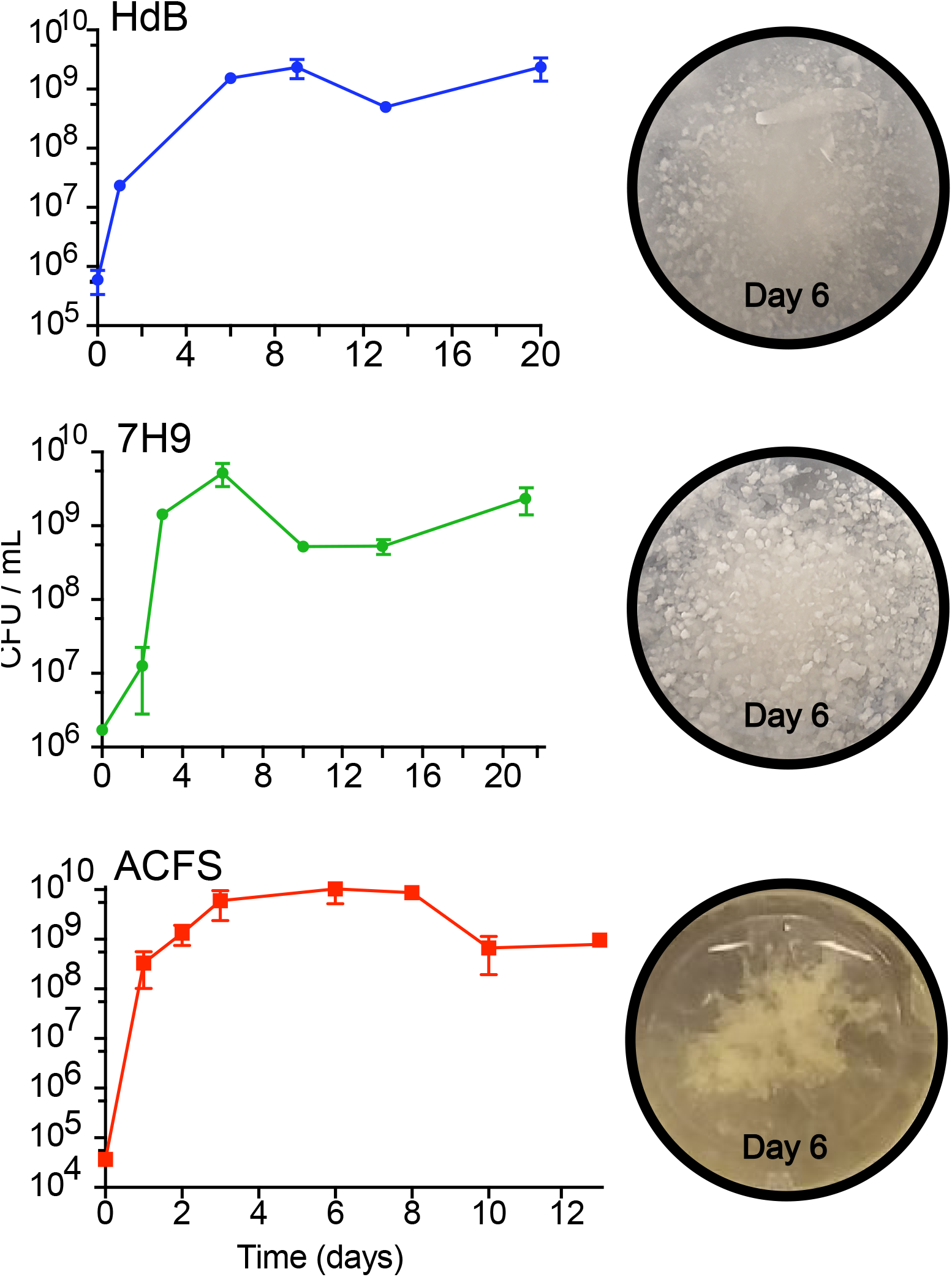
The development and macroscopic structure of *Mab* biofilms in different media. Colony forming units for biofilm-associated cells in standing culture over time. Images of corresponding wells taken at 6 days.

To probe the physiological differences between biofilm and planktonic cells in our assay, we tested their ability to transcriptionally and translationally respond to induction of a reporter. We transformed *Mab* with a plasmid that expresses Tag-RFP under the control of a constitutive P_GroEL_ promoter, and expresses eGFP under control of an inducible P_nitrile_ promoter. We allowed biofilms of this culture to mature for 6 days, then induced them with isovaleronitrile to induce expression of eGFP. After 24 hours of induction, we resuspended and fixed cells from the planktonic and biofilm populations in each media type, and visualized the cells by fluorescence microscopy (Fig. 3A). We found that in all media conditions, cells in biofilms are less likely than planktonic cells to express significant eGFP upon induction, compared to the constitutive Tag-RFP (Fig. 3B). This shows that biofilm cells are either less permeable to the inducer, or are less transcriptionally and translationally active than planktonic cells in the same culture well, or both. These data also validate our method of separating planktonic cells from biofilm cells, by showing separated cell types are physiologically distinct in all three media types.

**Figure 3.**
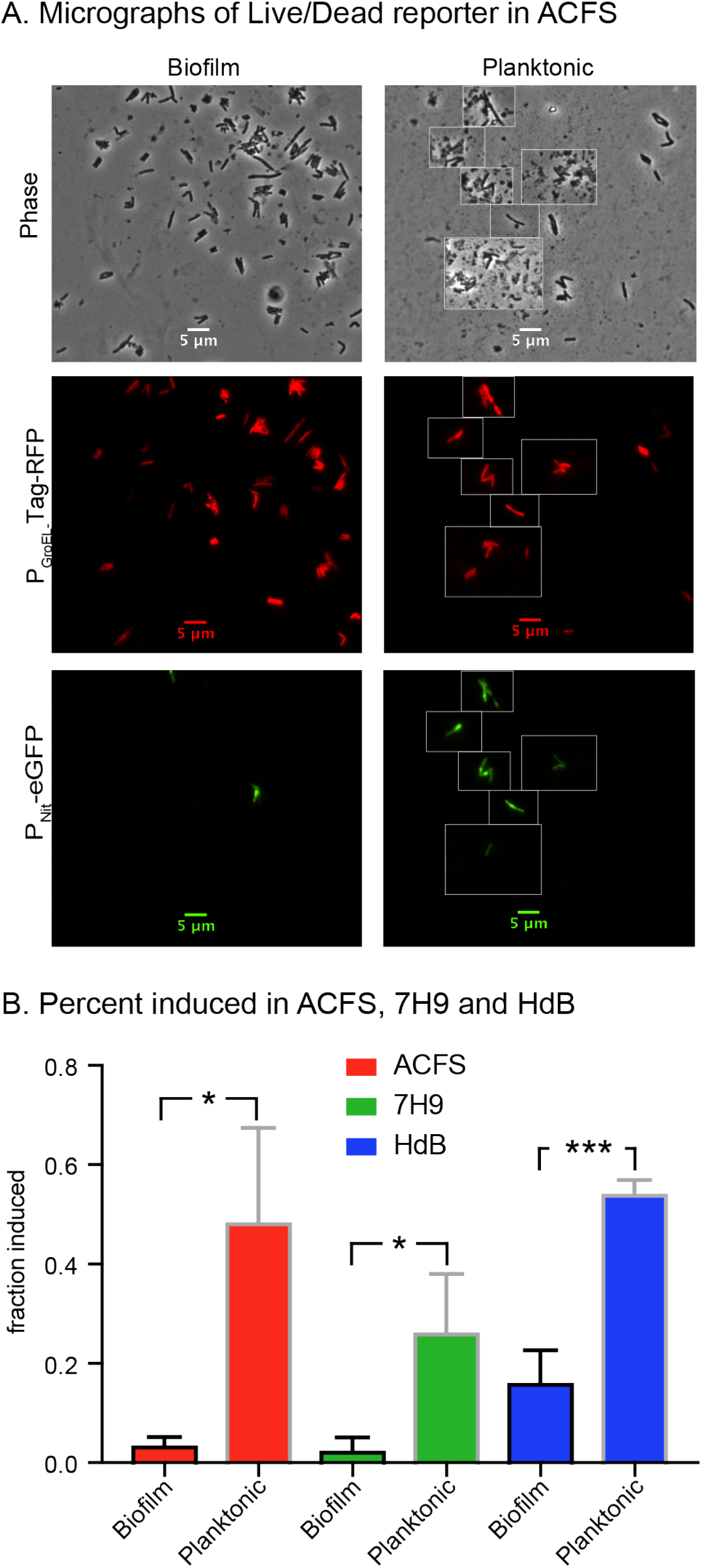
Cells in *Mab* biofilms are less responsive to a transcriptional inducer. (A) Micrographs of *Mab* cells carrying the pDE43-MEK-Nit-live/dead vector from biofilms grown for 6 days in ACFS, then induced with 10^-6^ isovaleronitrile for one day before fixation and imaging. Living cells that retain the vector show constitutive Tag-RFP signal, inducible cells also show eGFP signal. (B) Percentage of cells with average Tag-RFP signal intensities above 400 that also have average eGFP signal intensities above 100 in each condition. At least 100 cells each from three independent biological replicates were quantified for each condition. P-value of biofilm vs. planktonic for ACFS, 7H9 and HdB medias are 0.0152, 0.0271 and 0.0007, respectively. P-values were calculated using an unpaired t-test.

Because biofilm morphology varied so greatly between media types (Fig. 2), we hypothesized that media could affect glycolipid expression. We extracted surface glycolipids from planktonic smooth and rough strains grown in 7H9 and ACFS (Fig. 4A) as well as from mature biofilms grown in all three media conditions (Fig. 4B). We then separated the glycolipids by thin layer chromatography (TLC). TLC analysis shows that the smooth strain expresses similar levels and types of GPLs irrespective of media condition, while the rough strain produces minimal GPLs (Fig. 4A). In biofilms, the smooth strain expresses GPLs in all conditions (Fig. 4B,D). Based on previous GPL analyses (24), the purple spot near the top (Fig. 4B, spot c) is an unidentified wax, while the dark purple spot at the bottom (Fig. 4B, spot d) is trehalose dimycolate (TDM) (31). Both smooth and rough strains have increased TDM in ACFS compared to lab media, in both planktonic cultures (Fig. 4A) and biofilms (Fig. 4B). We observe a statistically significant increase in the wax (Fig. 4B,C) and TDM in ACFS (Fig. 4B,E). The levels of GPLs appear to be similar (Fig. 4B,D). These data show that media conditions affect the surface glycolipid expression of *Mab* and that *Mab* grown in ACFS media has higher levels TDM, a glycolipid that is a virulence factor in *Mtb* infections (22).

**Figure 4.**
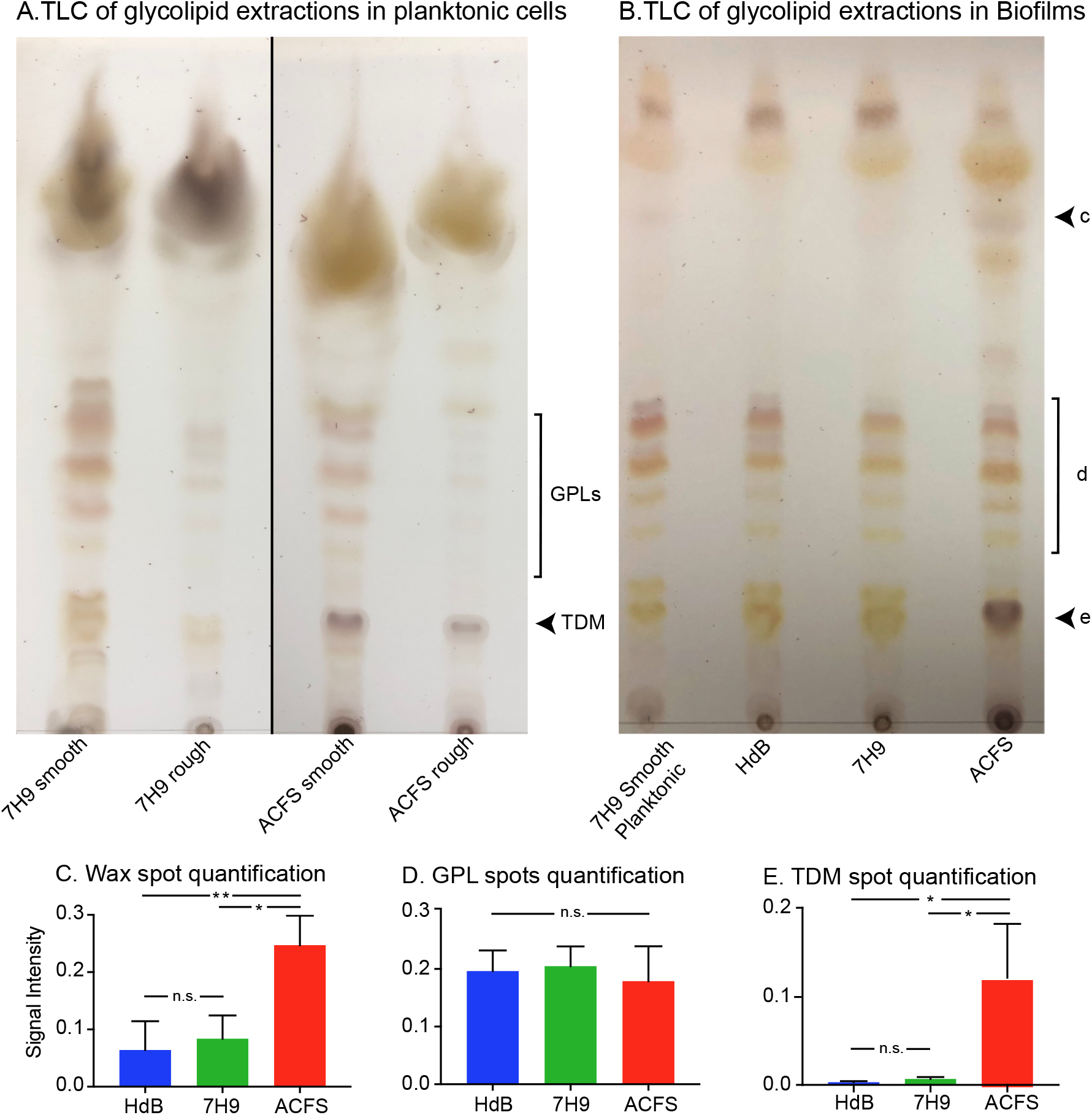
Glycolipid profiles are affected by media condition. (A) Thin Layer Chromatography (TLC) of glycolipid extractions from planktonic cultures of rough and smooth *Mab* strains grown in 7H9 and ACFS. GPLs = glycopeptidolipids, TDM= trehalose dimycolate. (B) Representative TLC of glycolipid extracts from mature biofilms of smooth *Mab* in Hdb, 7H9 and ACFS. Extraction from planktonic cells grown in 7H9 + Tween80 was used as a control sample. Black arrows indicate lipid species prominent in ACFS: c - unknown wax; d – GPLs; e - TDM. (C, D, E) Quantification of the signal intensity for each spot in (B) by means of spot densitometry. Values shown are the mean of three biological replicate experiments with error bars representing the standard deviation. (C) Wax, P-values between ACFS vs. Hdb and ACFS vs. 7H9 were 0.0079 and 0.0134, respectively. (D) GPLs, no significant differences. (E) TDM, P-values between ACFS vs. Hdb and ACFS vs. 7H9 were 0.0149 and 0.0171, respectively. P-values were calculated using one-way ANOVA with Tukey correction for multiple comparisons. Signal intensities for (D) were quantified by using the red channel only of the TLC images, to eliminate the signal from the yellow spots in the same area.

We sought to determine how biofilm maturity affects antibiotic tolerance. For these and all subsequent biofilm experiments, we compared survival of biofilm and planktonic cells from the same well of a 24-well plate for each replicate. We incubated *Mab* in ACFS media for 3, 6 or 11 days, then treated each well with two clinical drugs, clarithromycin and imipenem. We then measured survival over time and found that tolerance increases over time in standing culture, likely due to nutrient depletion of the media (Fig. 5AB). In 3-day and 6-day old biofilms, planktonic cells die off substantially upon treatment, while at 11 days both populations are quite tolerant (Fig 5A). 6-day old biofilms had higher tolerance compared to the planktonic cells in the same well at both 24 and 48 hours of treatment (Fig. 5B). From this, we conclude that at six days the biofilms in ACFS media are mature enough to substantially protect cells against antibiotics. Based on these results, and the fact that biofilms in all media types reached maximal cell density at six days (Fig. 2), we conducted all subsequent biofilm assays with 6-day old biofilms.

**Figure 5.**
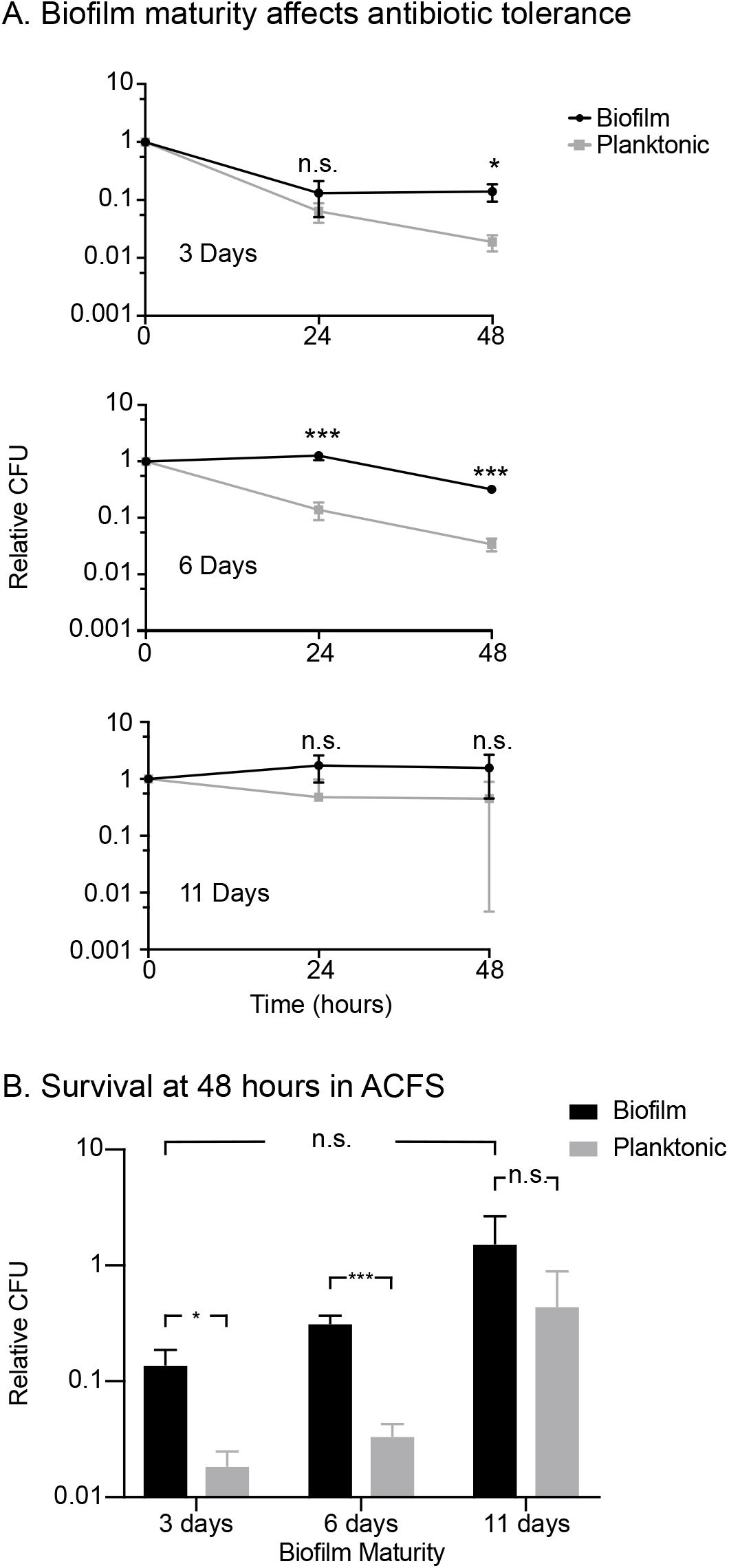
Antibiotic tolerance increases as ACFS biofilms mature. (A) Relative CFUs (CFU at each time point divided by the CFU value before treatment, *i.e*., t=0) from biofilms and planktonic cells from the same wells, grown for 3, 6 and 11 days in ACFS. Cultures were treated with clarithromycin (20 μg/mL) and imipenem (15 μg/mL) for 48 hours, and CFUs were measured at 0, 24 and 48 hours. (B) Survival from 48-hour CFU time points (triplicate wells). P-values between biofilm and planktonic cells are 0.01099, 0.0006, 0.1815, for 3, 6, and 11 days respectively. P-value for biofilms between 3 and 11 days is 0.09046. P-values were calculated using the student’s t-test.

In order to determine how media type affects the antibiotic tolerance of *Mab* biofilms, we treated 6-day old biofilms grown in different media types with varying concentrations of the clinically used antibiotics amikacin, clarithromycin and cefoxitin, and then plated both the biofilm and planktonic cells at 48 hours after treatment. We found that amikacin consistently killed planktonic cells at nearly an order of magnitude more than the biofilm cells from the same well, across media types. With clarithromycin and cefoxitin treatment, however, we saw much more variability. With both these antibiotics, planktonic cells and biofilm-associated cells were both highly antibiotic tolerant in both 7H9 and HdB. However, in ACFS, planktonic cells were much more sensitive to clarithromycin and cefoxitin than biofilm-associated cells. Clarithromycin was the only drug that was able to kill biofilm-associated cells at concentrations below 50 μg/ml, and this only occurred in ACFS media (Fig. 6).

**Figure 6.**
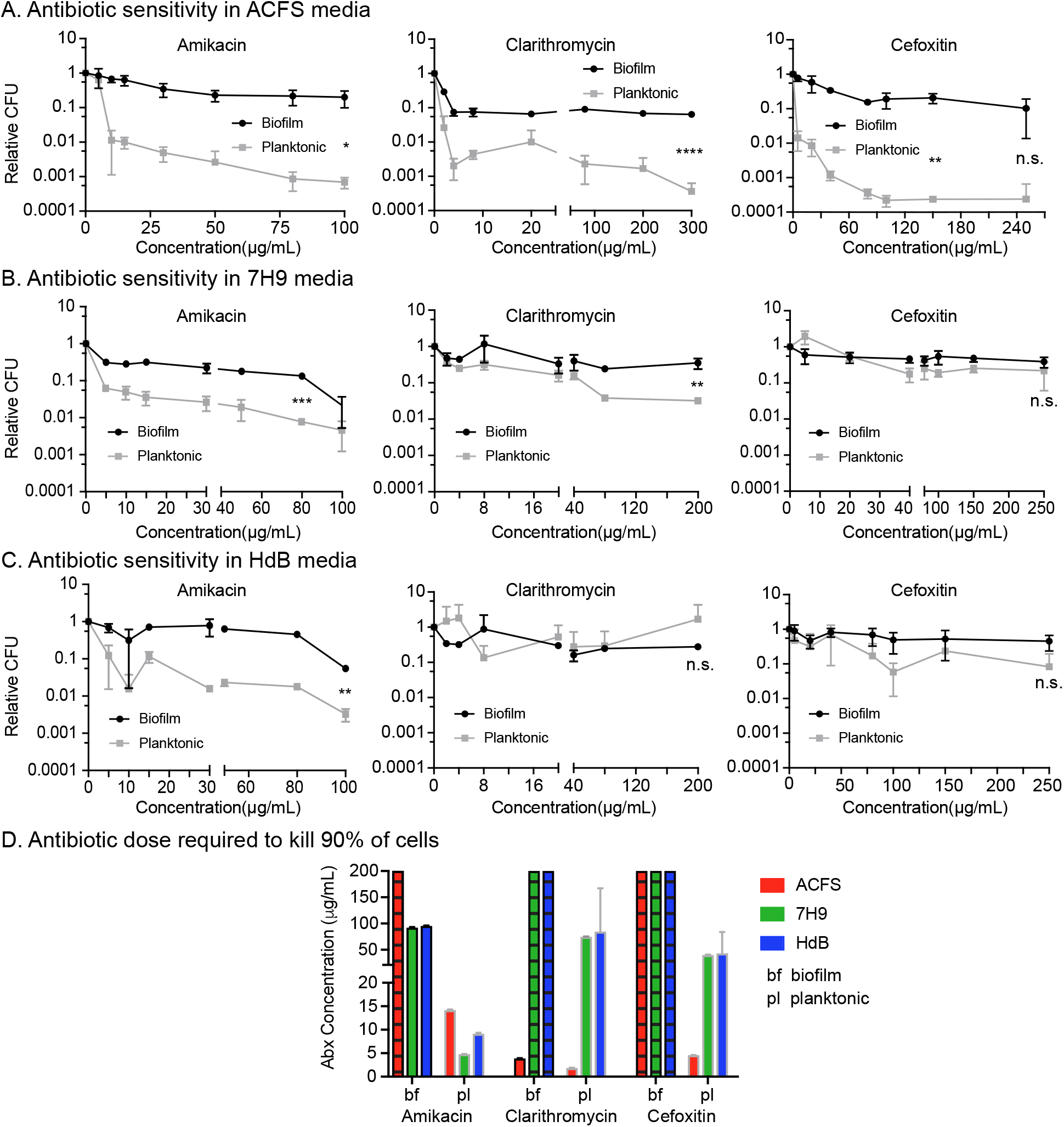
Antibiotic sensitivity varies by antibiotic and media type. (A, B, C) Relative CFUs of 6-day old biofilms and planktonic cells from the same wells after 48 hours of treatment with a range of antibiotic concentrations in 3 medias. Asterisks represent significance as measured by the student’s t-test; * = P ≤ 0.05; ** = P ≤ 0.01; *** = P ≤ 0.001; n.s. = P > 0.05. (D) LD90 (dose required to kill 90% of the cells) analysis of the data from A, B and C. Bars with hatched lines indicate that we were unable to observe killing of 90% of the population at any concentration used, so the LD90 is above 200 μg/mL. The LD90 was calculated for each replicate and the bars indicate mean and with standard deviation.

In our assays, the biofilm-associated cells are likely to have access to less oxygen than the planktonic cells. We sought to determine whether hypoxia could differentially affect cells depending on the media. We made cultures of *Mab* in tubes with minimal headspace, sealed the tubes with septum caps, and incubated them without shaking for 13 days. We then injected cefoxitin through the caps and measured survival after 48 hours. Results show that hypoxic *Mab* exhibits tolerance in both 7H9 or ACFS media (Fig. S2), comparable to that seen in biofilms (Fig.6AB). Thus, as for *Mtb* (32), hypoxia promotes antibiotic tolerance in *Mab* and is likely to be one factor in the antibiotic tolerance seen in biofilm-associated cells.

## Discussion

Bacterial physiology is enormously plastic. Bacterial cell size, shape, growth rates, surface properties and gene expression all change profoundly even between different lab media conditions (33-35). This plasticity makes it very difficult to determine the physiological state of a pathogen during infection. In this study, we sought to physiologically profile the emerging pathogen *M. abscessus* in two standard lab growth medias and one media designed to mimic the nutrient conditions in the lungs of Cystic Fibrosis patients (ACFS). We find that, compared to standard lab medias, ACFS media induces profound physiological changes in *Mab*, both in planktonic (Fig.1, Fig. 4) and biofilm states (Fig.4, Fig.6).

Bacterial biofilms are complex structures which, on the macro scale, vary physiologically according to developmental stage (36), surface type (37) and nutrient conditions (38). On the microscopic scale, there is also heterogeneity within a single biofilm: cells in different places within the biofilm express different surface factors and have different gene expression profiles (39-41). Thus, it is not meaningful to merely compare the phenotypes of “biofilm” and “planktonic” cells from a given species, as there is enormous phenotypic variety within each of these categories. In this work, we profiled some of the phenotypic diversity possible in *Mab* biofilms in order to assess the extent to which *in vitro* biofilm experiments might be relevant to the physiology of *Mab* biofilms within the lungs of CF patients. Our results show that *Mab* biofilms in ACFS media have important physiological differences from biofilms in the standard mycobacterial lab medias HdB and 7H9 (Fig. 2, Fig.4, Fig.6). The fact that such differences can be seen even between *in vitro* media conditions shows that *Mab* biofilm physiology is responsive to environmental conditions. We hypothesize that *Mab* physiology in HdB and 7H9 media is unlikely to represent its physiology during infection. Our results do not establish ACFS media as a proxy for infection, which of course includes many host immune factors that cannot be mimicked *in vitro*. However, these data show that nutrient conditions that might be encountered by *Mab* in the host could have an important role in the pathogen’s expression of virulence glycolipids and responsiveness to antibiotic treatment.

Even in experiments with the smooth *Mab*, which produces GPLs, we observed differences in the appearance of the biofilms that formed in the different media types. The biofilms in both HdB and 7H9 appeared flat and flaky, while the biofilms that formed in the ACFS media looked wet and formed tendrilly plumes (Fig. 2). These differences could be merely due to the chemical and viscosity differences between the media types, or they could be due to changes in *Mab* cell physiology in the different media types. We profiled the surface glycolipids and antibiotic sensitivity of cells in these different biofilms, and found that cells in ACFS biofilms are physiologically distinct: they display more trehalose dimycolate and a non-polar wax species than biofilms in 7H9 and HdB (Fig. 4) and they exhibit more relative antibiotic tolerance compared to planktonic cells (Fig. 6).

Because nutrient availability affects the physiology of cells within a biofilm, we sought to develop methods that would allow us to disentangle responses to nutrients from responses to biofilm formation. Other groups solved this problem by moving biofilms to fresh media each day (12, 19). We assumed that nutrients would not be unlimited during infection, and so worked to assess biofilm physiology in nutrient-limited culture (Fig. 2, Fig. 5). In order to control for the well-known effects of nutrient limitation on antibiotic tolerance (42), we compared biofilm and planktonic cells from the same culture wells. These cells are sharing the same depleted nutrients, so differences in physiology should be due to association with the biofilm, or to differences in access to oxygen. While we observed considerable antibiotic tolerance in 7H9 and HdB against clarithromycin and cefoxitin, there was similar tolerance in planktonic and biofilm-associated cells (Fig. 6B,C), even though the planktonic cells presumably have more access to oxygen at the top of the well. This implies that antibiotic tolerance in these media conditions is due to some other environmental factor, such as nutrient depletion, rather than due to the protection of the biofilm or to hypoxia. In the more nutrient-rich ACFS media however, we see much higher sensitivity to cefoxitin and clarithromycin in the planktonic cells, while the biofilm-associated cells are tolerant. Therefore, the physiological difference between biofilm and planktonic cells is greater in ACFS media than in typical lab medias.

Our results imply that the modest degree of antibiotic tolerance in biofilms that has been observed before (19) may be a function of the culture conditions. Our data show that mature biofilms grown in ACFS media afford considerable protection against three clinical antibiotics from different classes. While the ACFS media may mimic the nutrient conditions of sputum, it does not replicate a complex host environment with active immune cells and tissue structures, so the physiological state of *Mab* in a true infection remains unknown. However, our results imply, during infection in CF patients, *Mab* may express more of the virulence glycolipid TDM, and be more antibiotic tolerance that experiments in regular lab media might indicate. Thus, treatments that could help disrupt *Mab* biofilms or prevent their establishment could help clear these infections more quickly. A deeper understanding of the chemistry and genetics of biofilm formation and physiological responses in *M. abscessus* will be needed in order to determine the role of biofilms in infection and treatment recalcitrance and to develop better treatment protocols.

## Materials and Methods

### Media and culture conditions

All *M. abscessus* ATCC19977 cultures were started in 7H9 (Becton-Dickinson, Frankin Lakes, NJ) media with 5 g/l bovine serum albumin, 2g/l dextrose, 0.85 g/l NaCl, 0.003 g/l catalase, 0.2% glycerol and 0.05% Tween80 and shaken overnight at 37C until in log. phase. All *Mab* CFUs were performed by plating serial dilutions on LB Lennox agar. Three media types varying in nutrient richness were used for all biofilm assays: Tween80 was not added to any media for biofilm assay. Hartmans de Bont (HdB) Minimal Media was made as described (43), 7H9 for biofilms was made as above, without the Tween80.

### Artificial Cystic Fibrosis Sputum

Two protocols were combined in order to make our ACFS media (27, 44). In 500mL of ultrapure water, the unique components that comprise CF sputum were added as described (44); 5 g of porcine mucin (NBS Biologicals), 4 g of Deoxyribonucleic Acid (Spectrum Chemicals), 5.9 mg diethylene triamine pentaacetic acid (DTPA), 5 g NaCl, 2.2 g KCl, and 1.81 g Tris base. Amino acid stocks were prepared as described (27) and stored at 4C. Except for tryptophan, amino acids were added to the media. The ACFS media was adjusted to a pH of 7, filled-up to one liter, and autoclaved at 110C for 15 minutes. The media was cooled down at room temperature prior to adding tryptophan and 5 mL of egg yolk emulsion (Dalynn Biologicals). ACFS Media was stored at 4C.

### Biofilm assays

Log. phase biological replication cultures of *Mab* cultures in 7H9 were inoculated into tissue-culture treated 24-well plates with 2mL of the appropriate media to a final optical density of 0.02. Sufficient wells were inoculated in order to have biological triplicates for each time point or concentration in the assay. Prior to incubation, each plate was covered with a Breathe-Easy sealing membrane (Electron Microscopy Sciences) to allow air exchange and reduce evaporation. Standing biofilm plates were incubated at 37C. The Breathe-Easy membrane was removed after biofilm development in order to add antibiotics. The biofilm plates were then re-sealed and incubated again at 37C for 24 or 48 hours. For each data point, cells from biological replicate wells were used to plate for CFU calculations. To enumerate the planktonic cells, the supernatant was extracted from each well without disturbing the biofilm and plated. The remaining biofilm was then re-suspended in 7H9 + Tween80 and serial dilutions were plated and incubated at 37°C for 4 days. Serial dilutions were done in 96-well plates in HdB + Tween80 and plated on plain LB Agar plates. Relative CFU is the ratio between each CFU value and the initial CFU value at t=0 or concentration=0.

### Hypoxia Assay

*Mab* was grown in 7H9 until log-phase. In order to promote hypoxia, sterile glass tubes were filled near full (9 mL) with 7H9 and ACFS respectively. All tubes were inoculated to an optical density of 0.02, capped with a sterile rubber septum, and wrapped with parafilm to avoid uncapping. To establish growth cessation, standing tubes were incubated at 37C and CFUs were taken every couple days on biological replicates. Once we established that growth cessation happens at 13 days, new tubes were inoculated and incubated for 13 days. In order to avoid oxygenation, replicates were treated by injecting through the rubber septum with 80 μg/mL of cefoxitin. CFUs were taken upon addition of antibiotic (t=0), and 48 hours post-treatment. All time-points were plated on LB Agar and incubated for 4 days.

### Glycolipid extractions and Thin layer chromatography

Mature biofilms and planktonic cultures were used for glycolipid extractions. GPLs were extracted as described (34) and spotted on TLC Silica Gel 60 plates (Millipore Sigma). In short, cells were isolated and centrifuged. Pellets were resuspended for lipid extraction in 10mL chloroform/methanol (2:1). Extractions were done twice at room temperature for 24 hours and then centrifuged at 5000rpm for 30 minutes to collect organic supernatant. Organic samples (Lipids) were evaporated with nitrogen and resuspended in 1mL chloroform/methanol (2:1) and treated with equal volume 0.2M NaOH (in methanol). Lipids were incubated at 37C for 30 minutes and then neutralized with a few drops of glacial acetic acid. Solvents were evaporated and the lipids were resuspended with 4mL of choloroform, 2mL of methanol, and 1mL of water in each tube. Samples were mixed and then centrifuged at 5000 rpm for 10 minutes to collect the organic layer. The solvent was evaporated and the remaining lipids were resuspended in 100uL of chloroform/methanol (2:1) and spotted. TLCs were developed in chloroform/ methanol/ water (100:14:0.8) as described (24). Plates were visualized with 10% Sulfuric Acid (in ethanol) and baked for 20 minutes at 120C. The signal intensity of spots were analyzed and quantified using ImageJ software by means of spot densitometry. For spot d (TDM) the TLC image was split into single color channels and quantified by measuring the signal intensity from the red channel only. Each spot intensity value was normalized against all the spots in a lane (entire lane; all in triplicates). We re-ran the TLCs, when necessary, so that total lane spot intensities were similar between samples.

### Cell staining

For experiments on log. phase cells, *Mab* was stained with 3μL/mL of 10 mM NADA and incubated at room temperature for 15 minutes in growth media. Cells were then pelleted and resuspended in PBS+Tween80 and FM4-64 was added to a final concentration of 4 ng/mL and incubated at room temperature for 5 minutes. Cells were then fixed with 2% paraformaldehyde for one hour at room temperature, and washed and resuspended in PBS+Tween80 for imaging.

### Microscopy

*Mab* cells were resuspended as described for plating of planktonic and biofilm cells, then fixed with 4% paraformaldehyde in PBS at room temperature for 2 hours. Cells were immobilized on agarose pads and imaged using a Nikon Ti-2 widefield epifluorescence microscope with a Photometrics Prime 95B camera and a Plan Apo 100X, 1.45 NA objective. Green fluorescent images were taken using a filter cube with a 470/40 nm excitation filter and a 525/50nm emission filter. Red fluorescent images were taken using a filter cube with a 560/40nm excitation filter and a 630/70 emission filter. Images were captured using NIS Elements software and analyzed using FIJI and MicrobeJ (45).

