## Supplemental figures for "Biofilm-associated *Mycobacterium abscessus* cells have altered antibiotic tolerance and surface glycolipids in Artificial Cystic Fibrosis Sputum Media"

### Supplemental Materials

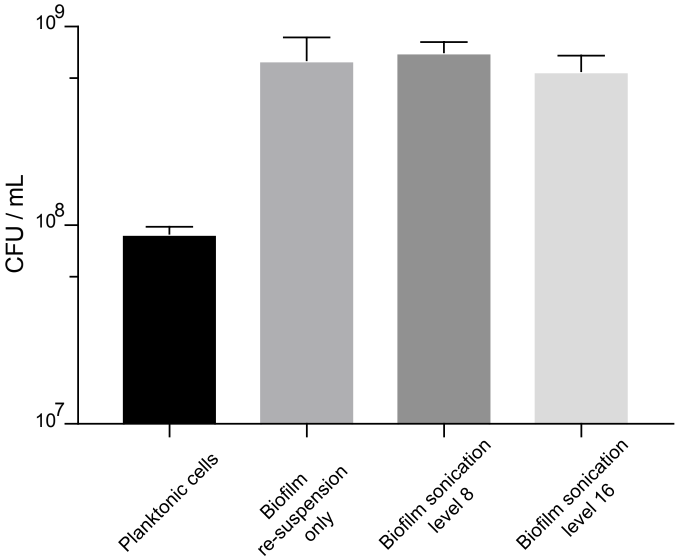


**Supplemental Figure 1. Mab Biofilm Sonication.** CFUs of mature biofilms after re-suspension and sonication. Planktonic cells were used as a control. Levels of sonication represent numbers on the amplitude control knob of a Microson Ultra Sonic Cell Disruptor XL.

*Hypoxia***.** We performed an assay in order to determine the extent to which hypoxia contributes to antibiotic tolerance in *Mab*. It is known that anaerobic conditions can induce antibiotic tolerance in *P.aeruginosa*, another common CF pathogen (1). Therefore, we aimed to compare the antibiotic tolerance in Mab planktonic cells subjected to anaerobic conditions, to antibiotic tolerance in mature biofilms at the bottom of our 24-well plates. We sealed planktonic cultures and left them stationary for 13 days, then treated them with cefoxitin and measured survival. We see in supplemental figure 2 that hypoxia does induce antibiotic tolerance in *Mab* when grown in both 7H9 and ACFS media. These data indicate that hypoxia could be one of the conditions that cause the biofilm-associated cells in our assay to be physiologically different from the planktonic cells. The hypoxic cells in ACFS are more sensitive to cefoxitin than those in 7H9 (student’s t-test; P-value 0.01344). This implies that either ACFS could retain more dissolved oxygen than 7H9 due to its greater viscosity, or that the nutrient conditions affect the way that cells respond to hypoxia.

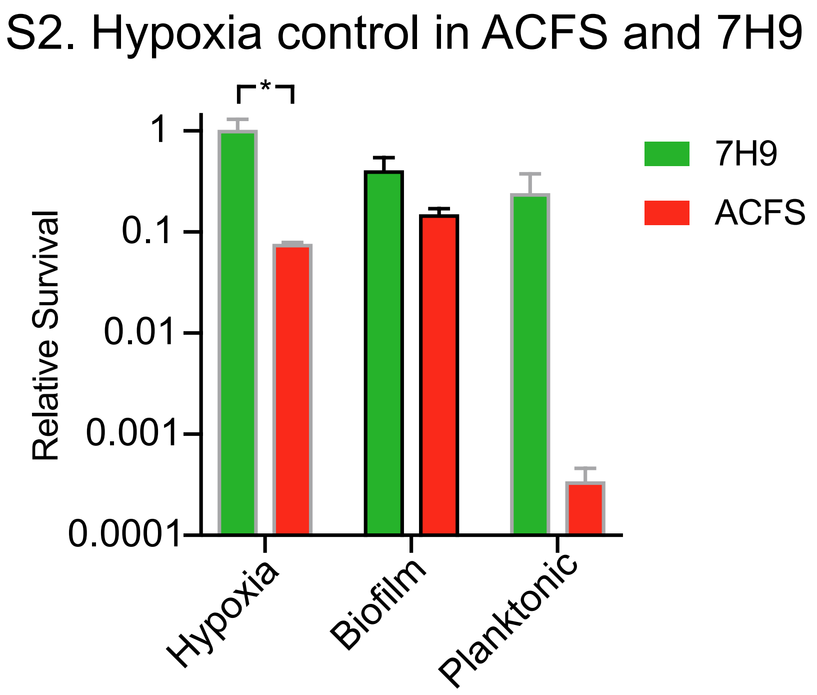


**Supplemental Figure 2. Antibiotic sensitivity of Hypoxic planktonic cells compared to antibiotic sensitivity in mature biofilms and planktonic cells.** CFUs of hypoxic cells 48 hours after treatment with 80ug/mL of cefoxitin (Hypoxia Assay). CFUs of mature biofilm and planktonic cells 48 hours after treatment with 80ug/mL cefoxitin (data from Fig. 6A, 6B).

*Antibiotic tolerance in the* Mab *rough morphotype***.** We isolated a “rough” morphotype in order to determine whether the media effects we observe on physiology are restricted to the smooth strains. In supplemental figure 3 (S3) we performed an antibiotic treatment course with mature biofilm and planktonic cells of rough *Mab*. Our original goal was to test the susceptibility of the pellicle, biofilm, and planktonic cells of rough *Mab*, and compare results to the smooth *Mab* data. Due to inconsistencies in pellicle formation within wells, and pellicle dispersion post-treatment, the pellicle data cannot be included. In addition, it must be noted that all results shown in S3 could be due to pellicle dispersal into the biofilm and planktonic populations. Despite dispersal, results do show that the trend in antibiotic susceptibility in smooth *Mab* holds in the rough *Mab* morphotype.

S3. Antibiotic sensitivity in Mab rough strain

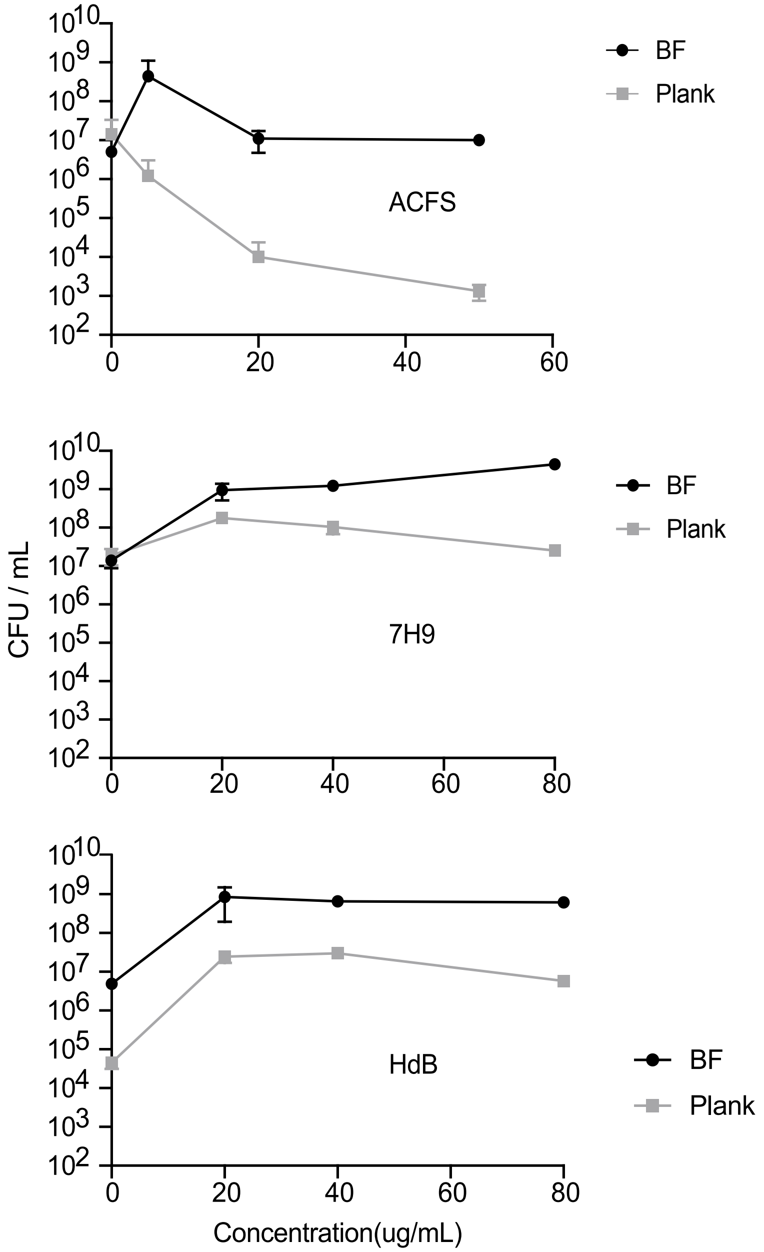


**Supplemental Figure 3. Rough strain exhibits comparable antibiotic susceptibility to smooth strain.** CFUs of 6-day old biofilms and planktonic cells of rough *Mab* treated with 80 μg/mL of cefoxitin for 48 hours. Compare to cefoxitin-treated smooth strain in all 3 medias (Fig. 6A, 6B, 6C). Pellicle dispersion post-treatment likely causes the increase in CFU between no drug and 20μg/mL.
